# Intracellular Regulation of a Serotonin-Gated Ion Channel Links Receptor Trafficking to Memory

**DOI:** 10.1101/2025.10.27.684756

**Authors:** Leona Cesar, Davide Zabeo, Emelie Aspholm, Dimitra Panagaki, Johanna Louise Höög, Julia Morud

**Author notes:** Corresponding author: Julia Morud, University of Gothenburg, Department of Chemistry and Molecular Biology, Box 462, 405 30, Gothenburg, Sweden.

## Abstract

Learning and memory arise from synaptic plasticity, the ability of neurons to modify connectivity through experience-dependent changes in receptor localisation and signalling. Here, we identify a short intracellular motif within the serotonin-gated ion channel LGC-50 that links molecular receptor dynamics to behavioural plasticity in *Caenorhabditis elegans*. Deletion of residues 363–379 in the intracellular M3–4 loop caused receptor clustering in intracellular compartments and abolished learning-induced redistribution without altering receptor function or immediate memory recall. Interestingly, animals expressing the truncated receptor failed to retrieve aversive memories one hour after training, revealing a role for receptor trafficking in memory stability. Combining molecular, ultrastructural and behavioural analyses *in vivo*, we show how intracellular receptor motifs govern experience-dependent plasticity. These findings demonstrate that precise receptor localisation and trafficking shape neural circuit adaptation and reveal a conserved mechanism by which receptor dynamics support the persistence and retrieval of memory across species.

## Introduction

Nervous systems possess an incredible capacity to adapt information processing over time, integrating sensory input with evolving internal states to enable experience-dependent decisions about when and how to act. This phenomenon, broadly referred to as neuronal plasticity, forms the basis for essential higher-order processes such as learning, memory formation and behavioural adaptation^1–3^. At the cellular level, plasticity is thought to emerge primarily from activity-dependent modifications in neuronal excitability within interconnected neural circuits^4,5^. These modifications can occur over a wide range of timescales, from rapid millisecond changes to long-lasting structural and molecular alterations that persist for days or longer^6^. A key mechanism driving such shifts in excitability involves the regulation of synaptic receptor abundance, particularly changes in the number and composition of ligand-gated ion channels (LGICs) at the postsynaptic membrane. Of special importance are the excitatory glutamate-gated receptors, including α-amino-3-hydroxy-5-methyl-4-isoxazolepropionic acid (AMPA) and N-methyl-D-aspartate (NMDA) receptors^7^, which mediate fast excitatory neurotransmission. In contrast the inhibitory gamma-aminobutyric acid A (GABA_A_) receptor provides a counterbalancing inhibitory influence on network activity^8^. By dynamically adjusting their abundance and distribution, these receptors recalibrate circuit responsiveness, enabling the brain to encode experience and adapt. The precise mechanisms governing these dynamic changes in receptor abundance remain poorly understood.

The trafficking of LGICs to the plasma membrane and their incorporation into synapses is tightly regulated through interactions with specific regulatory proteins^9–12^. Many of these mechanisms are activity-dependent, linking neuronal activation to plasma membrane trafficking, and thereby influencing processes such as memory formation^1,13,14^. Research to date has largely highlighted events proximal to the synapse, including plasma membrane trafficking of receptors, endosomal recycling and their anchoring by structural organiser proteins^10,15,16^. Within NMDA, AMPA and GABA_A_ receptors, distinct regulatory motifs have been identified as being critical for proper localisation to the plasma membrane and synapes^17–20^. However, the difficulty of tracking receptors *in vivo* means that dissecting these mechanisms at a molecular level in intact mammalian systems remains challenging. The nematode *Caenorhabditis elegans* (*C. elegans*) provides a powerful model, owing to its compact nervous system and lifelong transparency, which facilitate live imaging and genetic manipulation^21–23^. In addition, *C. elegans* carries genomic homologs of human LGICs as well as critical components of synaptic plasticity, such as scaffolding, anchoring, signalling and adhesion proteins, underscoring its utility as a model system for studying the molecular basis of synaptic plasticity^24–28^. Yet, the precise roles of these proteins in *C. elegans* and whether regulatory motifs function analogously to those in mammals is not well known.

*C. elegans* inhabits a complex microbial environment and must learn to discriminate between odours, associating them with either positive reinforcers, such as the presence of food, or negative ones, such as pathogenic bacteria^29–31^. These learning abilities allow short-term memory to be assessed in *C. elegans* by exposing animals to pathogenic bacteria, such as the *Pseudomonas aeruginosa* strain PA14, and subsequently testing their acquired avoidance of them^31,32^. Our recent work identified the homomeric cationic serotonin-gated channel LGC-50 as a critical component of serotonin-dependent pathogen avoidance learning^33^. A large intracellular domain in the loop between transmembrane helices 3 and 4 (the M3–4 loop) was shown to be important for plasma membrane localisation in *Xenopus laevis* oocytes expressing *lgc-50.* Truncation of residues 363-379 in the M3-4 loop (Figure 1A) increased the serotonin-induced current by approximately 1000-fold, suggesting enrichment of the channel on the plasma membrane^33^. In *C. elegans*, exposure of worms to PA14 bacteria caused LGC-50 to redistribute, increasing its intensity in the nerve ring. The receptor was found to play a critical role during signal integration in the interneuron RIA, underlying aversive associative learning to the pathogen PA14. However, the intracellular origin of this redistribution remains unclear and whether disruption of this process affects pathogen avoidance learning has yet to be determined.

**Figure 1.**
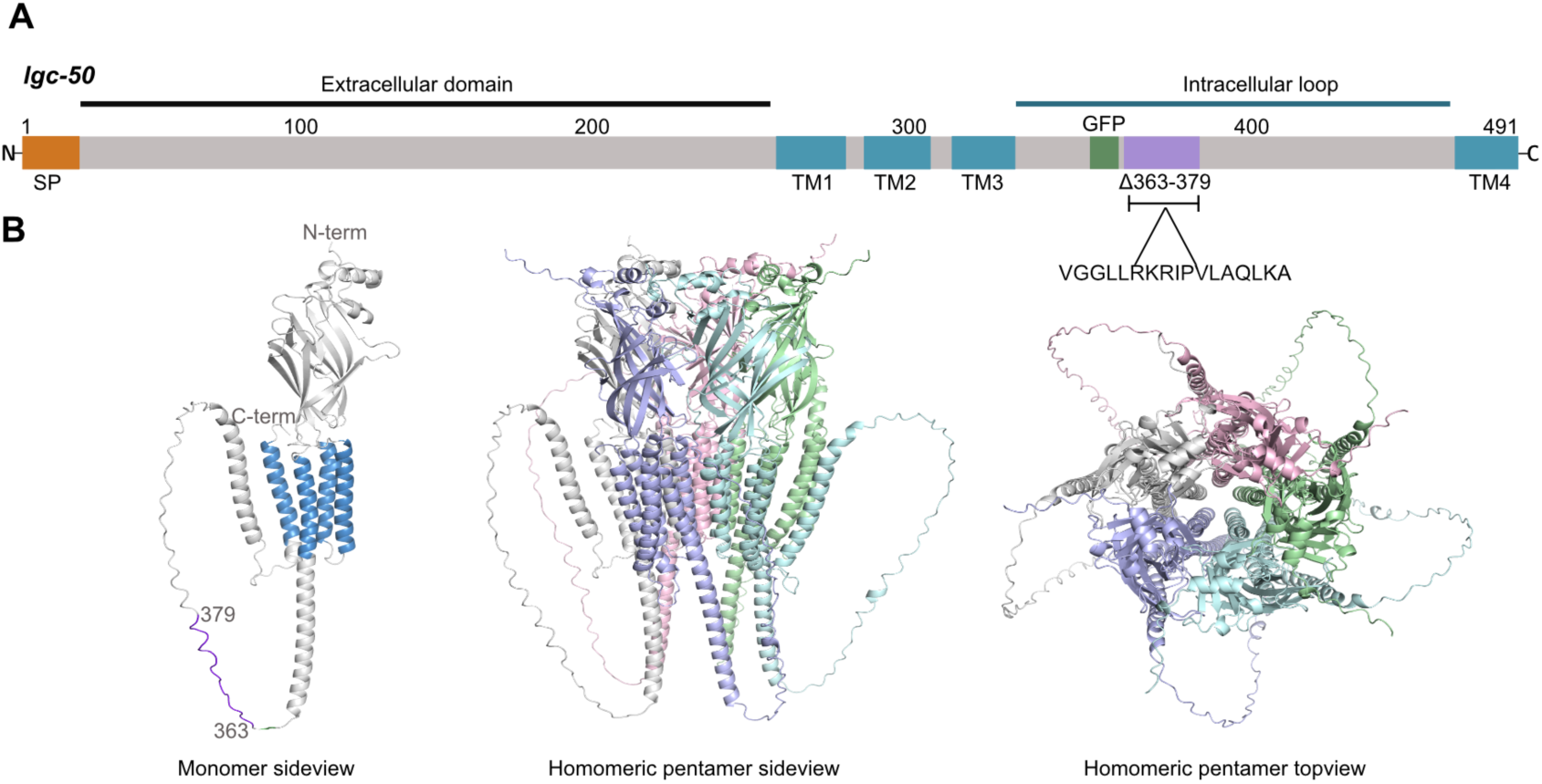
A. Graphical example of lgc-50 indicating location of the predicted signalling peptide (SP), CRISPR insertion site for GFP and the position of residues 363-379 in the transmembrane (TM) 3-4 intracellular loop. B. AlphaFold^34^ sideview and top view 3D models of an LGC-50 protein monomer and homomeric pentamer, with residues 363-379 highlighted in the monomer model (purple).

Here, we describe the function of the intracellular motif in LGC-50 that regulates the receptor’s subcellular localisation, using high-resolution electron and light microscopy. Truncation of this motif causes the receptor to form pronounced clusters at cellular membranes, independent of prior learning experience. Moreover, constant retention of the receptor at the plasma membrane interferes with activity-dependent trafficking, thereby impairing memory retrieval despite normal learning. This impairment is further reflected in the failure of the truncated receptor to undergo learning-induced changes in clustering, in contrast to the significant redistribution observed in control animals.

## Results

### An intracellular motif in LGC-50 regulates localisation and receptor clustering

Having previously observed that plasma membrane localisation of LGC-50 in a *Xenopus* heterologous system was influenced by a small truncation of the M3-4 loop (LGC-50Δ363–379)^33^, we sought to determine the effect of this truncation *in vivo*. For this purpose, we used a CRISPR edited strain with GFP inserted into the M3-4 loop of *lgc-50* (Figure 1A), this insertion site was previously validated to not interfere with LGC-50’s function^33^. In addition, we generated a deletion of residues 363-379 in *lgc-50* using CRISPR on the GFP background (Figure 1A-B).

To determine whether truncation of the receptor, LGC-50Δ363–379, causes alteration in the intracellular distribution or clustering of the protein in naïve animals, super-resolution light microscopy and ultrastructural examination with immuno-electron microscopy (immuno-EM) was employed.

We performed super-resolution microscopy in both the GFP-tagged *lgc-50* strain as well as the *lgc-50::GFPΔ363–379* strain (Figure 2A). Since the goal was to determine whether receptors were differently localised in areas relevant for aversive olfactory learning^31^ we focused our analysis on the nerve ring and the identification of fluorescent puncta. To enable unbiased quantification of fluorescent puncta, images were processed using a semi-automated image analysis pipeline^35,36^. The super-resolution fluorescence analysis revealed that the 363–379 truncation led to an increase in fluorescent puncta, both in total number and density (puncta/µm^2^), as compared to the full-length receptor (Figure 2B-E). Such accumulation in puncta could reflect a retention of the receptors at synaptic membranes. To ensure that this increase did not arise from higher overall expression of receptor in the LGC-50::GFPΔ363–379 strain, total fluorescence intensity within the nerve ring was measured with no significant difference between the two strains (Figure 2F).

**Figure 2.**
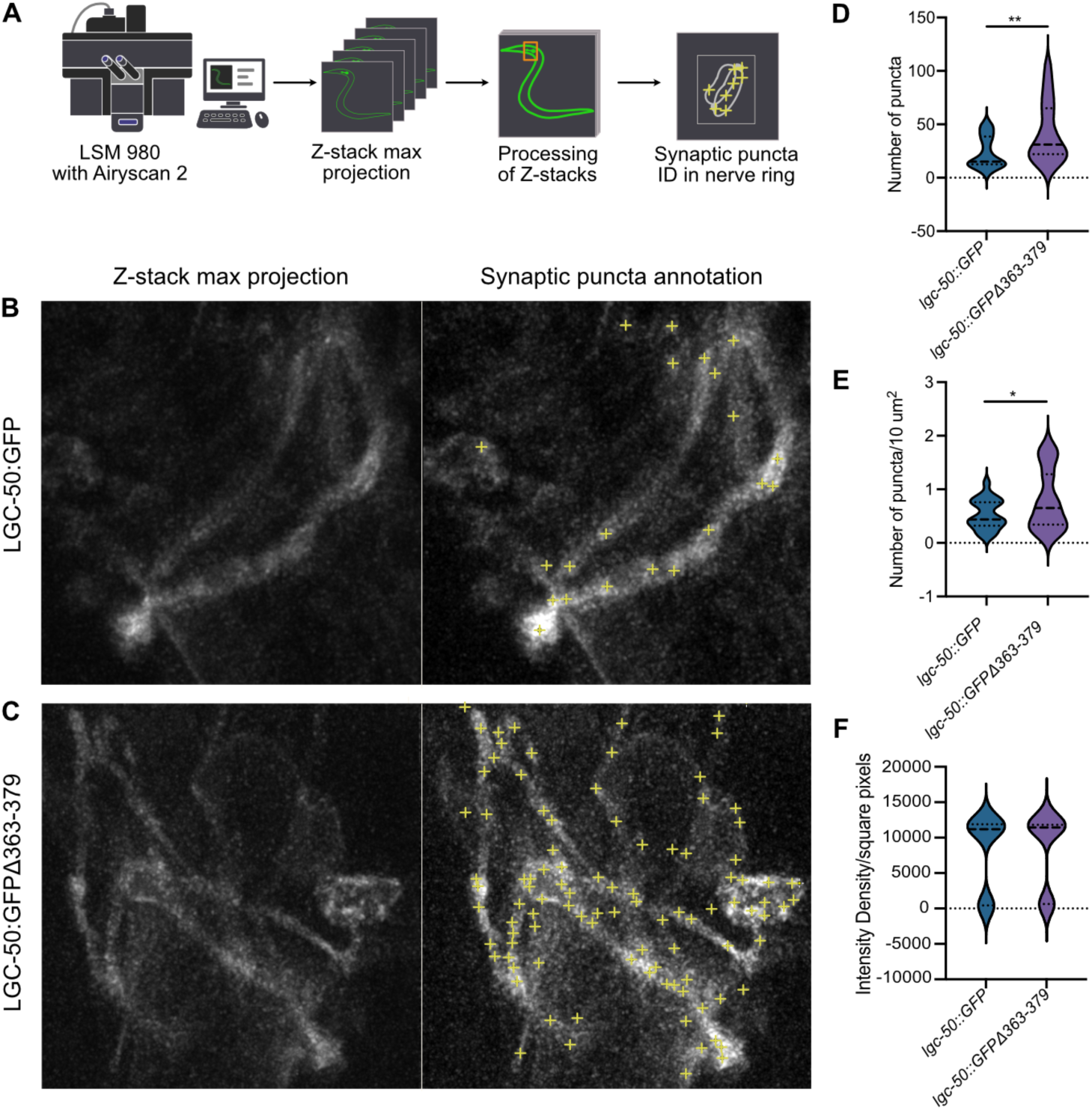
Deletion of residue 363-379 in LGC-50 significantly increases fluorescent puncta. A. Flowchart describing the work process for puncta quantification. B. Example image of GFP signal and puncta annotation in the nerve ring in full-length LGC-50::GFP(Ij120). C. Example image of GFP signal and the puncta annotation in the nerve ring in LGC-50::GFPΔ363–379(syb3377). D. Average total number of GFP puncta detected/worm in the nerve ring. E. Average number of puncta in nerve ring normalised by area/worm. F. Total fluorescent intensity detected in the nerve ring. N= 21 worms LGC-50::GFP, N= 19 worms LGC-50Δ363–379, presented as a violin plot with median, 1^st^ and 3^rd^ quartile marked with dashed lines, *P < 0.05 **P < 0.005, ns, no significant difference, calculated using paired t-test.

The increased puncta detected in this fluorescence analysis may reflect altered intracellular trafficking or more postsynaptic vesicular receptor pools^37^. As higher spatial resolution is needed to distinguish between these two scenarios, immuno-EM against GFP was applied ^38,39^. For optimal ultrastructural preservation, GFP-expressing worms were high-pressure frozen, embedded in resin and sectioned ultra-thin (Figure 3A). The GFP-tagged LGC-50 receptors, with or without truncation, were detected using gold-conjugated secondary antibodies. We compared the localisation and total number of gold particles per area across different tissues and cellular structures in the nerve ring of the worm (Supporting Table 1, Supporting Figure 1A). As *lgc-50* was previously shown to be expressed in both nervous and muscle tissue^33^ we focused our investigation on these tissues.

**Figure 3.**
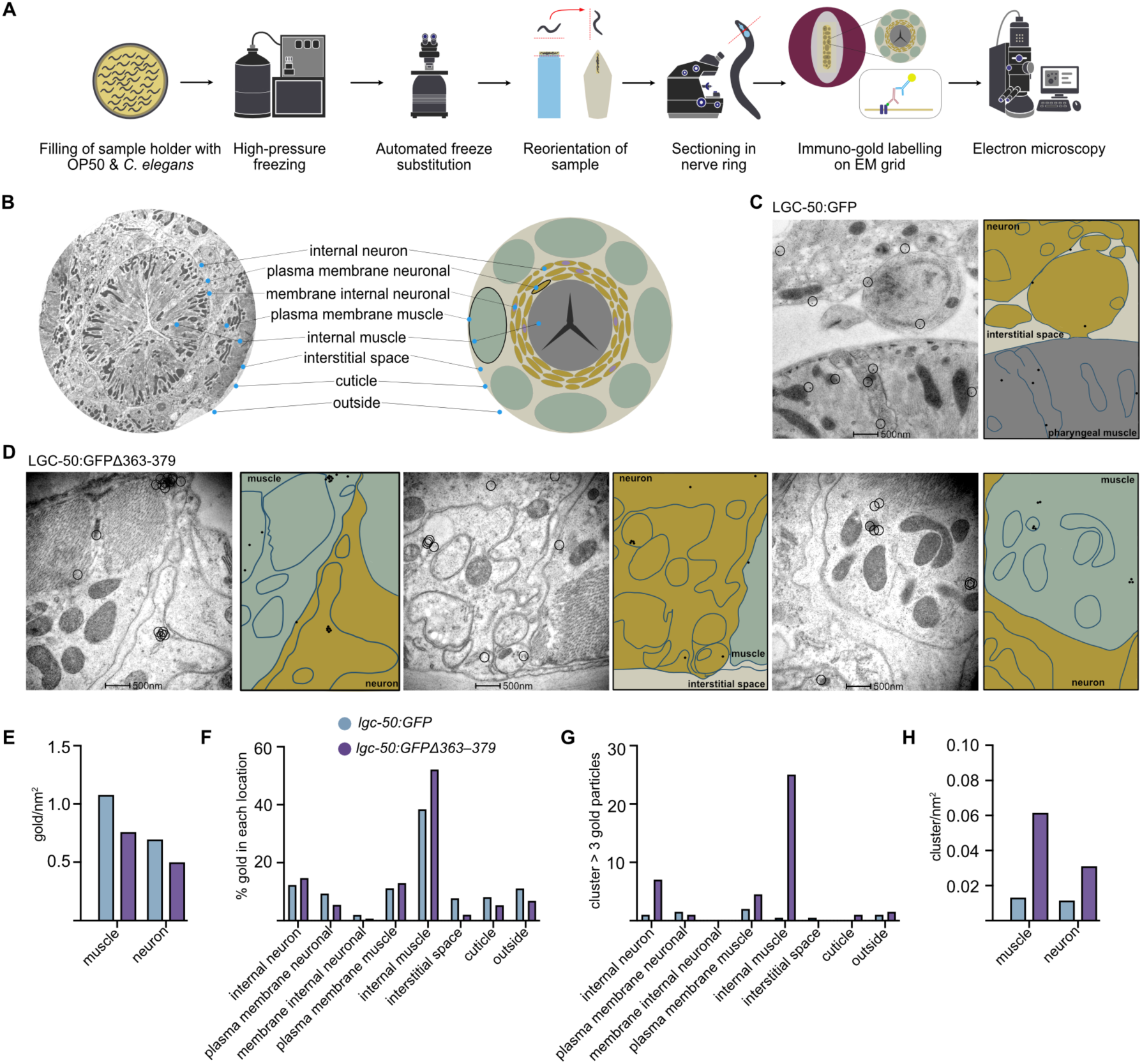
LGC-50::GFPΔ363–379 is localised in cluster in C. elegans. A. Graphical description of immuno-EM workflow. B. Cross section and cartoon of C. elegans cross-section through the nerve ring, describing where the different tissues and cellular structures quantified in the immuno-EM are identified. C. Micrographs and graphical description of gold-labelling in samples from full-length lgc-50::GFP(Ij120) animals. D. Micrographs and graphical description of gold-labelling in samples from truncated lgc-50::GFPΔ363–379(syb3377) animals that indicate high incidence of gold particles in clusters. E. Total number of gold particles per nm^2^ in muscles and neurons. F. Distribution of gold particles in different cellular locations. G. Number of clusters with more than three gold particles in different areas of the worm. H. Amount of clustered gold particles normalised by area. N=2 for each strain from which several micrographs were collected.

Interestingly, although the overall number of gold-labelled LGC-50 did not differ drastically between muscle or nervous tissue in the two strains (Figure 3H-I), the Δ363– 379 deletion strain displayed a markedly higher degree of gold clustering, with the gold particles more frequently appearing in groups of three or more in both neurons and muscles (Figure 3C-G, J-K, Supporting Figure 1B). In contrast, there were very few incidences of clusters with three or more gold particles in the full-length receptor samples (Figure 3C-D, J). In the Δ363–379 deletion strain, the most frequent neuronal localisation of GFP-labelled receptor clusters was in fact not at the plasma membrane, but rather in internal, non-membrane structures, possibly corresponding to endosomal or pre-synaptic vesicular compartments (Figure 3J-K). As a control, immuno-EM was also performed on wild-type *C. elegans* (N2) animals that did not express GFP. In these samples, very few gold particles were detected overall (Supporting Figure 1C), indicating high specificity of the antibody labelling for GFP. Although limited in its throughput (n=2 per strain), immuno-EM provides compelling high-spatial resolution evidence that this region of the M3-4 loop of LGC-50 plays a crucial role in its intracellular localisation.

### Truncation of residues 363–379 in LGC-50 do not interfere with aversive olfactory learning

It was previously reported that truncation of residues 363–379 in LGC-50 markedly increases the serotonin-induced current when the receptor is expressed heterologously in *Xenopus laevis* oocytes^33^, confirming that the truncated receptor remains functional. Given that *lgc-50(tm3712)* null mutant animals display defects in aversive olfactory learning^33^, we wondered whether the increased clustering observed for the truncated receptor might influence its function *in vivo* in a similar way.

To address this, we assessed the animals’ ability to perform chemotaxis towards both their standard food source *Escherichia coli* OP50 and the pathogenic bacteria PA14 (Figure 4). As a reference, we included the *lgc-50(tm3712)* null strain, which has previously been shown to exhibit normal chemotaxis towards OP50 but impaired aversive learning^33^. Both the *lgc-50* null and the *lgc-50::Δ363–379* truncated worms showed normal navigation towards OP50 and PA14, indicating intact basal chemotaxis (Figure 4A-F). To confirm that the day-to-day reliability was high, all behavioural experiments were performed over three individual days (Supporting Figure 2A-H).

**Figure 4.**
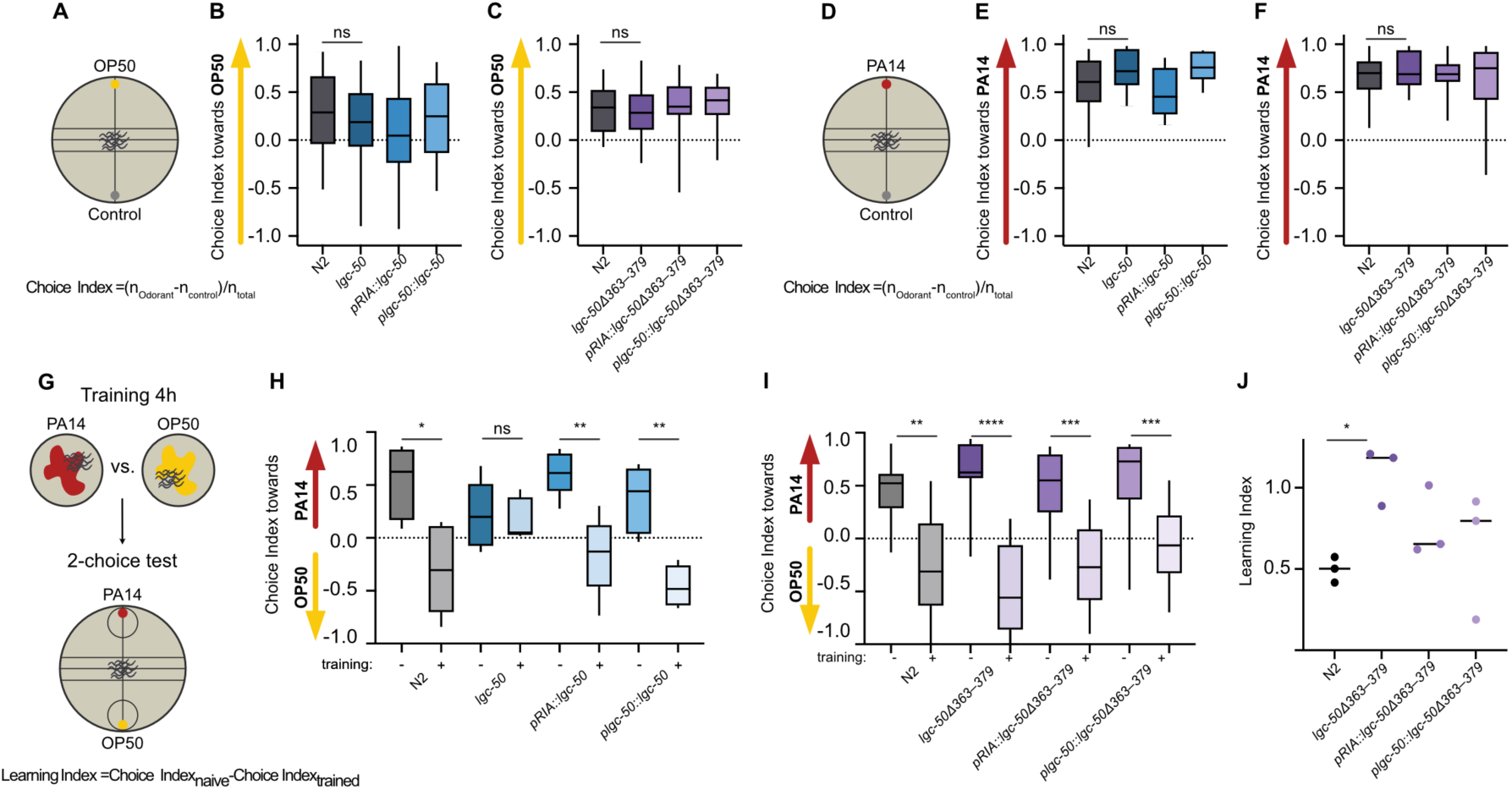
LGC-50Δ363–379 worms are capable of two-choice aversive olfactory learning. A. Schematic representation of experimental design. B-C. Chemotaxis towards OP50 for N2, lgc-50(tm3712), lgc-50(tm3712); pglr-6::lgc-50::SL2mKate2 referred to as pRIA::lgc-50, lgc-50Δ363– 379(syb3327), lgc-50(tm3712); plgc-50::lgc-50::SL2mKate2 lgc-50(tm3712); pglr-6::lgc-50Δ363–379::SL2mKate2 referred to as pRIA::lgc-50Δ363–379 and lgc-50(tm3712); plgc-50::lgc-50Δ363– 379::SL2mKate2. N = 17-30 plates/genotype. D. Schematic representation of experimental design. Chemotaxis towards PA14. N = 14-25 plates/genotype. G. Schematic representation of experimental design. H-I. Calculated choice indices after two-choice aversive olfactory learning towards PA14. N = 5 - 15 plates per genotype and learning condition. J. Learning index calculated from choice index after aversive olfactory learning. Each experiment was replicated over three days. *P < 0.05. **P < 0.01, ****P < 0.0001, ns, no significant difference, calculated by Welch’s t-test, data are shown as mean ± SEM.

Next, we tested the Δ363–379 strain’s ability to perform aversive olfactory learning, using the two-choice assay to measure olfactory preference to PA14 and OP50 after 4 hours of pre-exposure to PA14 (Figure 4G-J). Surprisingly, the *lgc-50::Δ363–379* worms performed significantly better than N2 controls, exhibiting roughly double the learning index score in this assay (Figure 4J). This suggests that the truncated receptor enhances the worms’ ability to remember the PA14 odour and associate it with negative experiences at this acute timepoint. As a control for assay conditions, we again included the *lgc-50(tm3712)* strain, which reproduced the expected learning-deficient phenotype, while expression of wild-type *lgc-50* in the RIA interneurons fully rescued the learning deficient phenotype (Figure 4H).

### Training with pathogenic exposure increases fluorescent puncta of LGC-50 in wild-type worms but not in the Δ363–379 truncated receptor

To further investigate whether receptor trafficking is altered by PA14 exposure we exposed animals to PA14 or OP50 for 4 hours and immediately imaged the nerve ring using super-resolution microscopy (Figure 5A). Quantification of fluorescent puncta was performed using the same semi-automated image analysis pipeline as previously described (Figure 5B-C).

**Figure 5.**
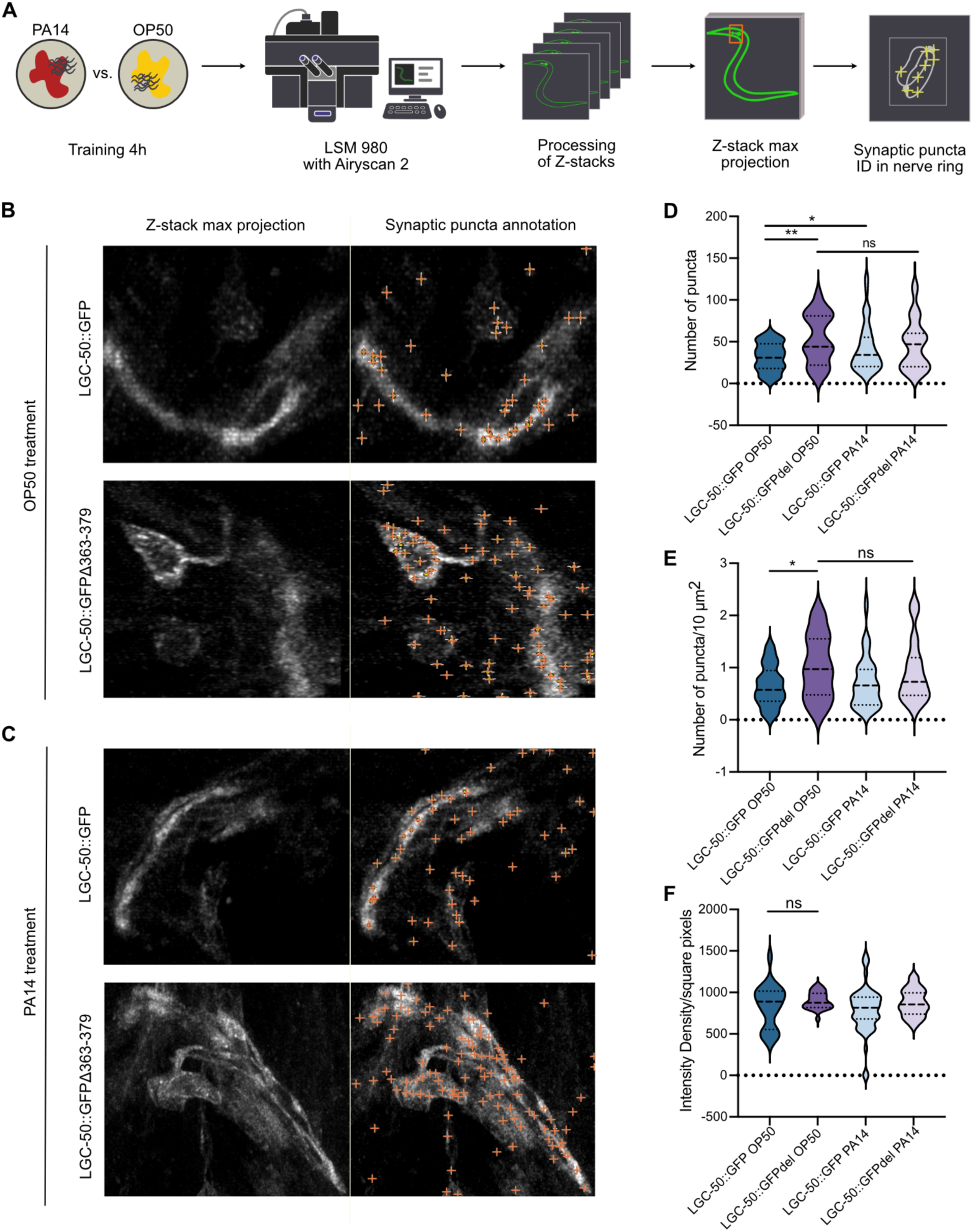
Exposure to PA14 for 4 hours increase fluorescent puncta of LGC-50::GFP(Ij120) in the nerve ring but not for TM3-4 loop truncated LGC-50(syb3377). A. Flowchart describing the work process for puncta quantification with PA14 exposure. B. Representative images of LGC-50::GFP worms with and without PA14 exposure. C. Representative images of LGC-50::GFPΔ363–379 worms with and without PA14 exposure. D. Average total number of detected puncta/worm with and without PA14 exposure. E. Average number of puncta normalised to area/worm with and without PA14 exposure. F. Total fluorescence measured in the nerve ring. N=29 worms LGC-50::GFP_OP50_, N=19 worms LGC-50::GFP_PA14,_ N=51 LGC-50Δ363–379_OP50_, N=30 worms LGC-50Δ363–379_PA14_, , presented as a violin plot with median, 1^st^ and 3^rd^ quartile marked with dashed lines, *P < 0.05 **P < 0.01, ns, no significant difference, calculated using an unpaired t-test.

Consistent with our earlier findings, the Δ363–379 truncated strain exhibited a significantly higher number and density of fluorescent puncta compared to wild-type animals (Figure 5D-E). However, in contrast to wild-type worms, in which PA14 exposure induced a robust increase in puncta number relative to OP50 exposure, the Δ363–379 strain showed no detectable change in puncta count or density following training (Figure 5D-E). Importantly, total fluorescence intensity did not differ between the two strains nor in the same strain before and after PA14 exposure (Figure 5F), indicating that the observed differences reflect receptor redistribution rather than altered protein expression levels. Together, these results demonstrate that pathogen-induced learning triggers a redistribution of LGC-50 in wild-type neurons, whereas the Δ363–379 truncation abolishes this plastic response. This provides further evidence that the intracellular motif deleted in the truncated receptor might play a crucial role in activity-dependent receptor trafficking and synaptic plasticity.

### LGC-50Δ363-379 are unable to retrieve memory already after 1 hour

To further explore if we could manipulate LGC-50 membrane trafficking, we expressed *lgc-50* without its predicted N-terminus signalling peptide^40^ (Δ2-19) on the *lgc-50(tm3712)* deletion background in the interneuron RIA (Figure 1). We asked if this would influence the aversive olfactory learning towards PA14 by restricting the receptor from accessing the endoplasmic reticulum (ER) membrane and subsequently inclusion into the plasma membrane via the secretory pathway. Surprisingly, the worms expressing the signalling truncated form of *lgc-50* were able to learn to discriminate between PA14 and OP50 after a 4-hour training period to the same degree as N2 control worms (Figure 6A-B). This suggests that LGC-50 might use an alternative signal-anchor domain to substitute the N-terminus signalling peptide. Possibly, the first transmembrane helix (M1) possesses hydrophobic residues that might be sufficient for ER membrane insertion. This process may involve the endoplasmic reticulum membrane complex (EMC), as has been reported for similar LGICs^41^.

**Figure 6.**
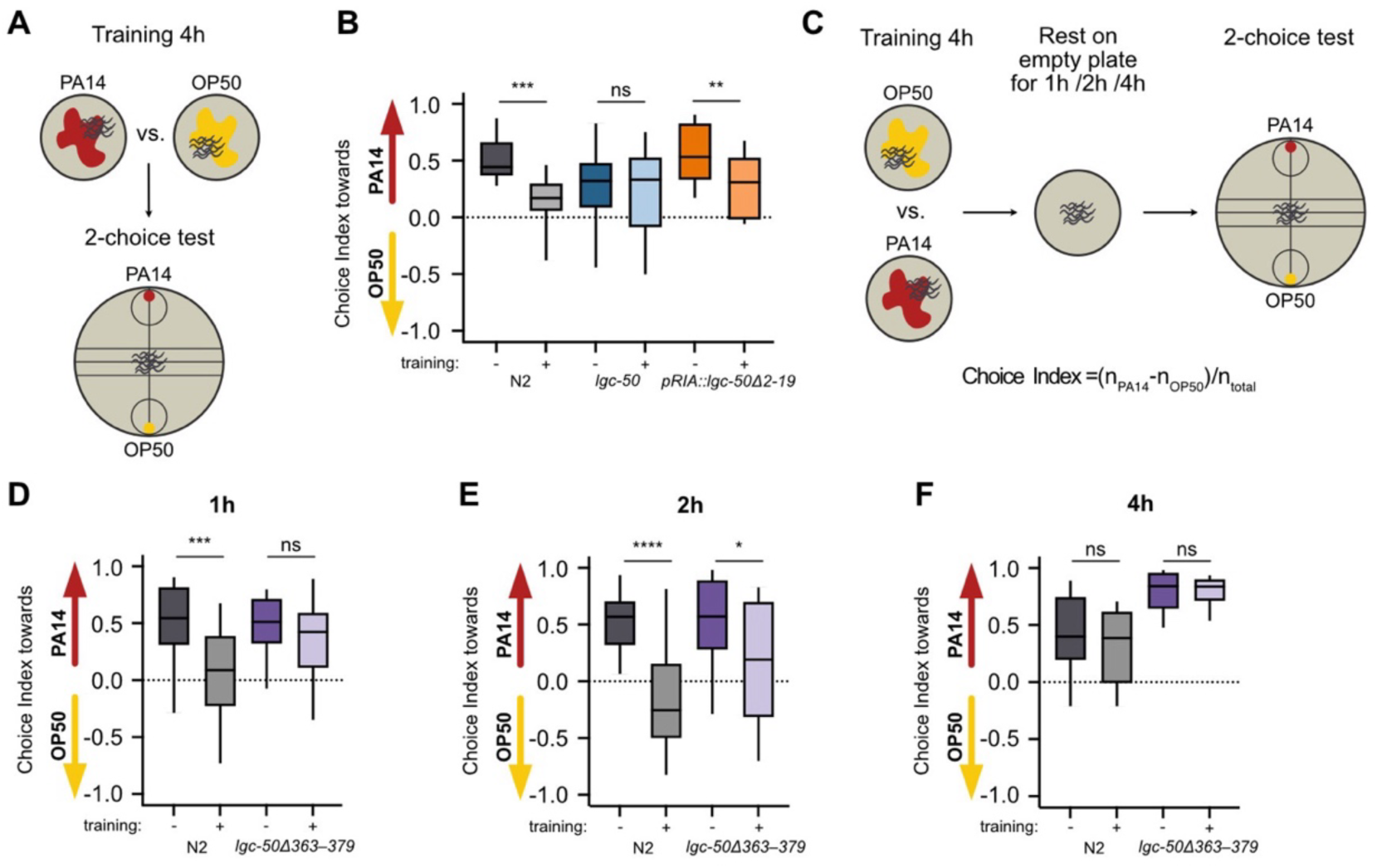
LGC-50Δ363–379 has impaired memory retrieval ability. A. Schematic representation of experimental design two-choice assay. B. Calculated choice indices after two-choice aversive olfactory learning towards PA14 for N2, lgc-50(tm3712) and lgc-50(tm3712); pglr-6::lgc-50Δ2-19::SL2mKate2 referred to as pRIA::lgc-50Δ2-19. N = 14-15 plates per genotype and learning condition. C. Schematic representation of experimental design for two-choice assay with delayed memory test. D-F. Choice indices after 1, 2 or 4 hours for N2 and lgc-50Δ363– 379(syb3327). N = 5-20 plates per genotype, learning condition and timepoint. Each experiment was replicated over three days. *P < 0.05. ***P < 0.001, ****P < 0.0001, ns, no significant difference, calculated by Welch’s t-test, data are shown as mean ± SEM.

Given that our initial learning data suggested that the *lgc-50::Δ363–379* strain exhibited normal or even enhanced aversive learning, we next assessed its memory retention over time, as a measure of synaptic strength and memory stability^42,43^ (Figure 6C). Animals were pre-exposed to PA14 for 4 hours, then allowed to recover on OP50 plates for 1, 2 and 4 hours before being tested in the two-choice olfactory assay. Consistent with previous reports^44^, N2 worms retained the learned aversion and successfully discriminated between the two odours for up to 2 hours after training (Figure 6D-F). In contrast, the *lgc-50::Δ363–379* worms had already lost the memory at the 1-hour time point (Figure 6D-F), corresponding to one hour of recovery between the end of the PA14 pre-exposure and the start of the two-choice testing.

This finding indicates that, while the Δ363–379 truncation does not impair the acquisition of aversive learning, it causes a rapid decay in memory retrieval, suggesting a defect in synaptic consolidation or maintenance of memory-related plasticity^45^. One possible explanation is that the truncation disrupts learning-induced receptor trafficking of the receptor or its recruitment to the membrane, thereby compromising the reinforcement of synaptic connections necessary for sustained memory retention. Our finding provides a compelling demonstration of how molecular changes in receptor localisation can manifest as measurable alterations in behaviour, offering direct evidence between synaptic plasticity mechanisms and memory expression in a living organism.

### Interaction partners to LGC-50 includes proteins of importance for synaptic strength and for the secretory pathway

The intracellular M3-4 loop of various LGICs has been repeatedly implicated in receptor localisation, membrane trafficking and protein complex assembly, both through interactions with other proteins^46^, and via distinct amino acid motifs within the loop itself^20,47^. Based on our imaging and behavioural data, the 17-residue segment of the M3-4 loop in LGC-50 (Figure 1) appears to be critical for membrane receptor trafficking and the stabilisation of synaptic connections with implications for memory retrieval. To identify potential binding partners to the LGC-50 interactome that could mediate these processes, we cultured large volumes of GFP-tagged *C. elegans* expressing either full-length or Δ363–379 truncated LGC-50 protein. Proteins were immunoprecipitated using anti-GFP magnetic beads and subsequently analysed by liquid chromatography-mass spectrometry (LC-MS) (Figure 7A). Both strains yielded a comparable number of interacting proteins (Supporting Figure 3), though differences were observed in the abundance and identity of specific binding partners (Figure 7B-D). The total count of peptide spectrum matches (PSM) were duplicate corrected and compared between the two genotypes. Several proteins of interest were found to be interacting with LGC-50 (highlighted in Figure 7B), such as proteins reported to be part of a multipass membrane protein machinery (NRA-2, nicalin) that is involved in the insertion and folding at the endoplasmatic reticulum^48^. Furthermore, UNC-116 was identified as a potential protein partner. UNC-116 has been shown to be involved in modulating synaptic strength through its involvement in rapid delivery, repositioning, and removal of receptors to the plasma membrane^49^. We also identified the copine protein NRA-1 that has previously been reported to interact with LEV-1^50^, a *C. elegans* nicotinic acetylcholine receptor. Copine proteins are known for being involved in directing proteins to membranes upon Ca^2+^ binding^51^, which could fit with an activity-dependent membrane trafficking model for LGC-50. To build hypothesises on the interactome of LGC-50 and protein partners that could be of importance for regulating its membrane localisation we specifically focused our analysis on proteins that could be of importance for ion channels and receptors, or for the secretory pathway. We compared the differential interaction between the full-length protein and the loop deletion (Figure 7C-D) and found several interesting proteins in these two classes to be differently pulled down with either of the protein versions. Together, these data suggest that LGC-50 interacts with components of both the synaptic and secretory machinery, highlighting a potential molecular framework through which intracellular receptor partners coordinate trafficking, localisation and ultimately, memory-related plasticity.

**Figure 7.**
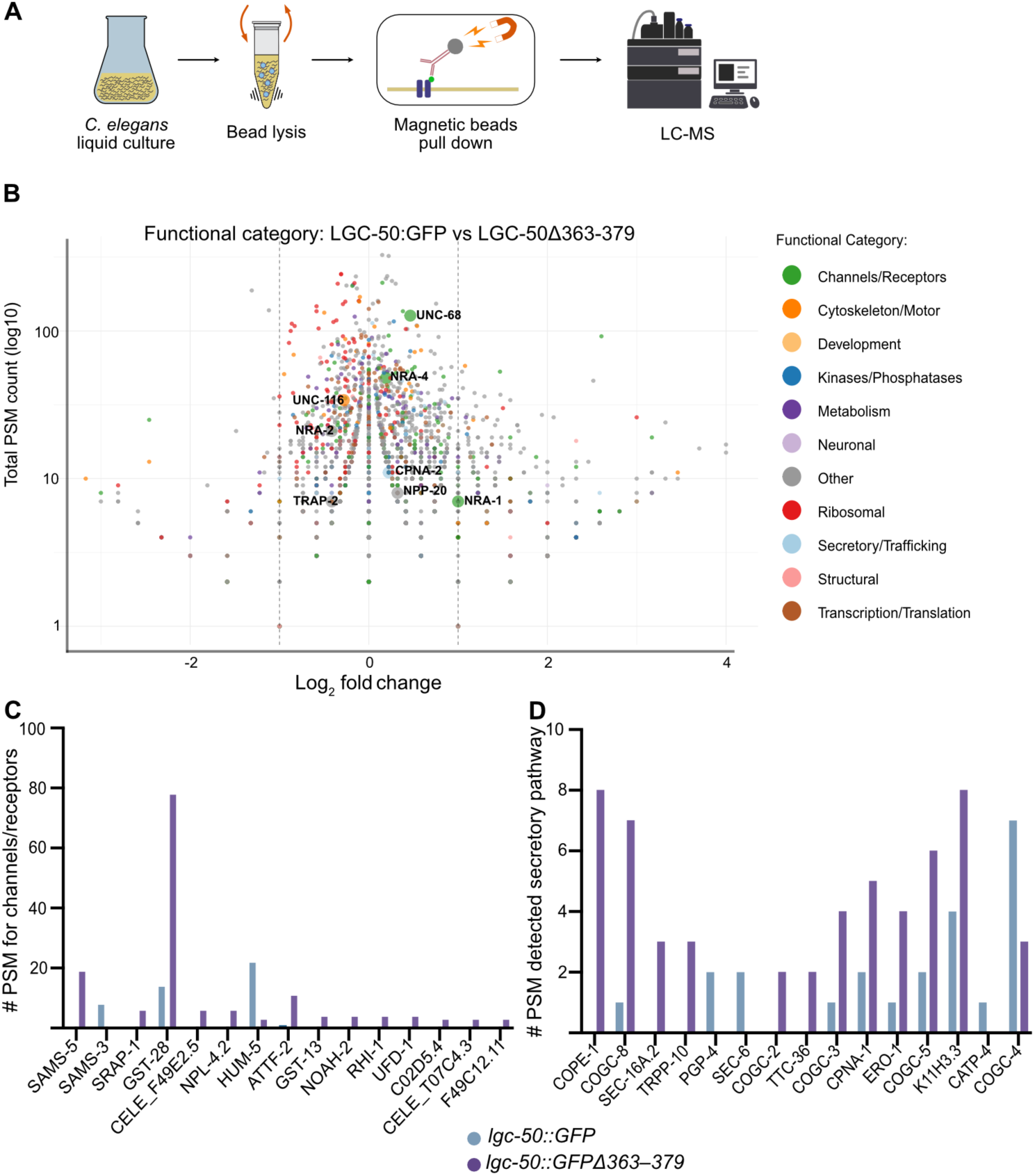
Immunoprecipitation with LC-MS analysis of LGC-50::GFP(Ij120) and LGC-50::GFPΔ363–379(syb3377). A. Workflow for immunoprecipitation experiments in C. elegans. B. Log_2_ fold change calculation of total PSM count. C. Quantitative comparison of the proteins that differ the most between genotypes for the channel-or receptor-relevant proteins. Key proteins of biological interest with significant fold changes are highlighted in the volcano plot. D. Quantitative comparison of the proteins that differ the most between genotypes for secretory-relevant proteins. Analysis was performed with pseudocount to handle zero values. Significance threshold was set to ±1.0 log_2_ units (corresponds to ∼2-fold change). Due to single replicate design, traditional statistical significance testing was not performed.

## Discussion

Neural circuits continuously adapt their function in response to experience, a process known as synaptic plasticity, which underlies learning and memory formation. At the cellular level, plasticity often arises from activity-dependent redistribution of neurotransmitter receptors at the synapse. In this study, we identify a short intracellular motif within the M3–4 loop of the serotonin-gated ion channel LGC-50 as a critical determinant of receptor trafficking and memory retrieval in *C. elegans*. By combining electron and super-resolution light microscopy, behavioural assays and proteomic analysis, we show that deletion of residues 363–379 alters receptor localisation, abolishes learning-induced redistribution, and leads to impaired memory retrieval without affecting initial learning.

Our imaging data demonstrate that deletion of residues 363–379 in LGC-50 leads to increased receptor clustering and a pronounced accumulation of receptors in intracellular compartments, likely representing endosomal or pre-synaptic vesicular pools. Despite normal overall expression levels, these receptors fail to redistribute following pathogen exposure, in contrast to wild-type animals, which display a clear increase in receptor puncta after learning. The behavioural consequences of this mislocalisation are notable as the truncated receptor supports normal or even enhanced learning but cannot sustain memory retrieval over time. Together, these findings indicate that this short intracellular motif is necessary for activity-dependent receptor trafficking and stabilisation of memory-related synaptic changes. Many Cys-loop LGICs are targeted to the endoplasmic reticulum (ER) and inserted into membranes through internal hydrophobic segments rather than classical cleavable signal peptides. The first transmembrane region (M1) has been shown to function as a signal-anchor, that could enable ER targeting and correct membrane insertion independent of a signalling peptide^41^. Consistent with this, our finding that deletion of the predicted N-terminal signal sequence did not disrupt LGC-50 function or learning suggests that the receptor likely uses an internal hydrophobic segment to mediate membrane targeting. However, further work will be needed to confirm the precise sequence elements responsible for LGC-50 insertion and topology in the ER membrane.

Our data highlight the M3–4 intracellular loop as a critical regulatory region for receptor localisation and trafficking. This loop is known to mediate protein–protein interactions, quality control of assembly and trafficking signals across multiple ion channel ^52,53^. Within this region, specific short motifs often act as endoplasmic reticulum (ER) retention or export signals. Arginine-based retention motifs such as RxR or RKR have been shown to prevent ER exit for various channels, including K_ATP_ and GABA_B_ receptors, until masked by the correct subunit^47,54^. The deleted segment in LGC-50 contains an RKR sequence, suggesting that this region may function as a conditional retention motif regulating receptor export from the ER. Removal of this motif could lead to premature or unregulated trafficking, resulting in accumulation of receptors in intracellular compartments. Additionally, the deleted segment includes serine and threonine residues, which could serve as phosphorylation sites influencing trafficking or receptor mobility. Similar regulatory mechanisms are well described for AMPA-type glutamate receptors, where phosphorylation of GluA1 by PKC or PKA promotes surface insertion during long-term potentiation^55,56^. Whether analogous phosphorylation-dependent mechanisms exist for LGC-50 remains to be determined, but such regulation could explain the activity-dependent redistribution observed in wild-type animals.

The trafficking pattern we observe for LGC-50 resembles activity-dependent receptor dynamics seen in mammalian systems. In hippocampal neurons, AMPA receptors are inserted into the plasma membrane during learning through the conventional secretory pathway, then move laterally to synapses^57,58^. Distinct AMPA receptor subunits display different trafficking behaviours, for example, GluA1 rapidly exits the ER and reaches the plasma membrane^59^, whereas GluA2 is more tightly regulated and often retained in intracellular puncta^60,61^. These differential dynamics fine-tune synaptic strength and contribute to the persistence of memory traces. Analogously, LGC-50 undergoes learning-induced redistribution to synaptic regions in response to exposure to the pathogen *Pseudomonas aeruginosa* PA14. The failure of the Δ363–379 receptor to respond to this training suggests that the deleted motif is essential for coupling receptor localisation to experience-dependent signalling. Thus, LGC-50 provides a tractable model to link molecular trafficking events to behavioural plasticity, allowing visualisation of receptor dynamics *in vivo* at synaptic resolution. Behaviourally, the Δ363–379 truncation separates learning from memory retention. Worms expressing the truncated receptor can acquire an aversive association but rapidly lose it within an hour, indicating that while synaptic signalling may initially be intact, synaptic stabilisation mechanisms are compromised. This parallels mechanisms observed in vertebrate systems, where maintenance of potentiated synapses depends on sustained receptor insertion, retention and recycling at the postsynaptic membrane^47,62^. The inability of LGC-50Δ363– 379 to redistribute after learning might thus prevent reinforcement and the strengthening of synaptic connections, leading to a failure to reactivate the relevant neuronal circuits during memory retrieval.

In wild-type animals, exposure to PA14 resulted in a marked increase in LGC-50 puncta, suggesting that learning triggers a redistribution of the receptor into discrete intracellular or synaptic compartments. An important question is whether these puncta persist for as long as the learned aversive memory is maintained and whether they subsequently dissipate once the memory fades. Such a temporal correspondence would support the idea that the observed puncta reflect the dynamic flow of receptors between intracellular pools and synaptic membranes, thereby mirroring the molecular consolidation and decay of memory. The intracellular localisation of these puncta may also indicate association with endosomal or vesicular compartments, consistent with receptor trafficking through recycling or sorting pathways rather than stable accumulation at the plasma membrane. If so, the increased puncta observed after PA14 exposure could represent a transient accumulation of receptors in trafficking intermediates that accompany activity-dependent redistribution to the synapse. In contrast, the absence of such regulation in the Δ363–379 mutant might indicate that the truncated receptor is trapped within these compartments and unable to complete the trafficking cycle required for synaptic insertion. This scenario would explain why receptor clustering is elevated but static in the mutant and supports a model in which the 17– amino acid intracellular motif enables the dynamic receptor cycling necessary for maintaining memory-related synaptic plasticity.

Our proteomic analysis identified several proteins that differentially associate with full-length versus truncated LGC-50. Among these were components of the endoplasmic reticulum membrane complex (EMC), such as NRA-2/nicalin, implicated in the insertion and folding of multi-pass membrane proteins^48^, and UNC-116, a kinesin motor protein involved in vesicular transport and synaptic receptor delivery^49^. These interactions suggest that LGC-50 trafficking is coordinated with cellular machinery that regulates protein folding, transport, and synaptic positioning. Disruption of these interactions could underlie the altered localisation observed for the truncated receptor.

Although the precise mechanism by which LGC-50 enters the secretory pathway remains unresolved, our findings indicate that regulated receptor trafficking is required for maintaining, but not acquiring, learned behaviour. The redistribution of LGC-50 after pathogenic exposure likely reflects a dynamic trafficking event linked to learning. The intracellular puncta observed may represent transient endosomal or vesicular pools involved in receptor recycling rather than stable membrane insertion. We propose that, after learning, LGC-50 cycles through these compartments toward synaptic sites, whereas in the Δ363–379 mutant, receptors become trapped in these intermediates, preventing the trafficking needed for memory maintenance.

By directly linking receptor trafficking to behavioural plasticity, this study reveals how molecular events at the level of individual receptors translate into network-level memory stability. More broadly, our findings provide a mechanistic framework for understanding how activity-dependent trafficking of neurotransmitter receptors underpins the persistence and retrieval of memory, a principle likely conserved across nervous systems from nematodes to mammals.

## Methods

### Resource availability

#### Lead contact

Further information and requests for *C. elegans* strains and plasmids is to be sent to and will be fulfilled by the lead contact Julia Morud Lekholm

### Materials availability

Materials generated in this study, including strains, plasmids and clones, are freely available from the lead contact upon reasonable request.

### C. elegans

Unless indicated otherwise, *C. elegans* worms were cultured at 20 °C on nematode growth medium (NGM), plates were seeded with *Escherichia coli* OP50. Transgenic rescue strains were produced by microinjecting plasmid DNA into the gonads of young adult hermaphrodites, and progeny carrying stable extrachromosomal arrays were subsequently selected. Offsprings with stable array transmission were selected for isolated strains. The following alleles were generated by SunyBiotech (Fuzhou, China) using CRISPR/Cas9-based genome editing: lgc-50::Δ363–379(syb3327) and lgc-50::GFP::Δ363–379(syb3377). CRISPR generated lines were backcrossed to our laboratory stock of wild-type worms (N2) four generations before being used.

### Molecular Biology

*C. elegans* gDNA, cDNA or promotor regions were amplified out of N2 DNA libraries using reverse transcription PCR with either Q5 polymerase (NEB, MA, USA) or Phusion and Phire polymerases (ThermoFisher Scientific). Gene fragments were assembled using the HiFi assembly protocol (NEB, MA, USA) or In-Fusion Snap assembly protocol (Takara, CA, USA) into the pDESTR4R3II vector. Unless specified, the promotor sequences used were approximately 2 kb upstream of the start codon.

### Electron microscopy

#### Sample preparation for Electron Microscopy (EM)

Day-1 adult worms were grown to high densities on NGM plates that had been generously seeded. The 0.1 mm cavity of the A type specimen carrier (Wohlwend GmbH, Switzerland) was filled with *C. elegans* and OP50 acting as a cryoprotectant^64^ using a worm pick. A Type B freezing carrier (Wohlwend GmbH, Germany) was dipped in hexadecene and placed flat side down on top of the carrier. The carriers were then frozen in a high-pressure freezer Compact 3 (Wohlwend GmbH, Switzerland) and transferred directly to the automatic freeze substitution (AFS) unit, (Leica, Germany). The samples were substituted with 2% uranyl acetate (Electron Microscopy Sciences, USA) dissolved in 10% methanol and 90% acetone^66,68^, but with the adaptation that the uranyl acetate was left on the sample overnight at -90°C. The samples were then washed in acetone before stepwise infiltration at -50 °C with 20%, 40%, 50%, 80% and 100% freshly prepared Lowicryl HM20 resin (Polyscience, USA) diluted in acetone, 2 hours per step. Infiltration continued overnight with 100% Lowicryl HM20 resin. UV polymerisation was initiated at -50 °C and continued during a controlled temperature rise to room temperature. Polymerisation proceeded for five days at room temperature. To reorient the sample, the top part of the sample cylinder containing the worms was cut off with a sawed, and the edges were trimmed. The sample was then placed in a flat embedding mould in the desired orientation and re-embedded in Agar 100 (Agar Scientific, UK). The samples were cut into 70 nm ultra-thin sections using a diamond knife (Diatome, Switzerland) fitted to a Ultracut E microtome (Leica, Germany) and placed on a copper slot grid (PELCO, 1 x 2mm, 3.0mm O.D.) (Ted Pella, USA) coated with 1% formvar (TAAB, UK) in chloroform.

### Immuno-EM

Thin sections were post-fixed in 1% paraformaldehyde in phosphate buffered saline (PBS) for 10 minutes, then washed 3 x 1 minute in PBS. The grids were then blocked for 1 hour in 0.1% fish skin gelatine (Merck, Germany) and 0.8% bovine serum albumin (BSA) in PBS. Immunolabelling was carried out by applying a 1:10 dilution in blocking solution of a rabbit anti-GFP antibody (Abcam, UK) overnight at 4 °C, followed by a three 20-minute in PBS. Then a 1-hour incubation at room temperature with a 1:20 dilution of a 10 nm gold-conjugated goat anti-rabbit secondary antibody (Aurion, Netherlands) was done. After another three 20-minute washes in PBS, the sections were postfixed with 2.5% glutaraldehyde (Thermo Fisher Scientific, USA) for 1 hour. Following 3 x 1 minute washes in ddH_2_O, the grids were imaged.

### EM image Acquisition

Samples were imaged on a Tecnai T12 transmission electron microscope (Thermo Fisher Scientific, USA) equipped with a Ceta 16M CMOS camera (Thermo Fisher Scientific, USA).

### Light Microscopy

The worms were immobilised on 2% agarose pads using a solution of 75 mM sodium azide in M9 buffer. Imaging was performed using a Zeiss LSM 980 with an Airyscan 2 detector and a 40× water-immersion objective (Zeiss, Germany). For trained samples, worms were exposed to *Pseudomonas aeruginosa* PA14 or OP50 training plates for 4 hours under the same conditions as in the behavioural assay before imaging.

### Chemotaxis

The preparation of plates and worm preparation and handling for the chemotaxis assay was done as previously described^63^, with some modifications. Fresh overnight liquid cultures of PA14 and OP50 were prepared from single colonies the day before the experiment. The following morning, the cultures were diluted and regrown until they reached OD600 of 1. For the chemotaxis assays, 25 µL of the PA14 or OP50 cultures (OD600 = 1) were spotted onto the assay plates and paired with 25 µl of lysogeny broth (LB) medium as a control. The plates were used in the assay within max. two hours after preparation. The day-1 adult hermaphrodites (∼50) were washed from the plate with M9 and transferred to the assay plates. After running the assay for 1h 15µl of sodium azide was dropped on the bacteria spots and the plates were stored at 4°C.

### Aversive olfactory training and learning

The aversive training with the pathogenic bacterium PA14 was performed as previously described^30,33,65^, with some modifications. To prepare the training plates, 100 µl of PA14 or OP50 culture was spotted onto NGM agar and incubated at 25 °C for two days. For the day of the experiment fresh overnight liquid cultures of PA14 and OP50 were prepared from single colonies. In the morning, the cultures were diluted and regrown until they reached OD_600_ of 1. For the learning assays, 25 µl of the PA14 and OP50 cultures (OD_600_ = 1) were spotted onto opposite sides of the assay plates. The assay plates were incubated at room temperature during the training period or during worm washing and were always used within max. two hours after preparation. Worm handling, washing and transfer to the assay plates followed previously described procedures^63^, except that the assays were performed on NGM plates rather than chemotaxis (CTX) plates. On the day of the experiment ∼70 day-1 adult hermaphrodites were transferred to PA14 or OP50 plates and exposed for four hours at room temperature before testing. The worms were washed from the training plates with M9 and transferred to the assay plates. After 1h 15 µl of sodium azide was dropped on the bacteria spots and the plates were stored at 4°C. For quantification, only worms within a 25 mm radius around each bacterial spot were included. For the time series memory retrieval assay, worms were washed from the training or control plates and placed on an empty plate with a small amount of OP50 during the incubation time before the test, after which they were washed off the plate and tested in the two-choice assay as described above.

### Immunoprecipitation from *C. elegans*

Large-scale worm cultures were prepared in liquid media, as previously described^67^. Synchronised worm populations were grown in S medium supplemented with high concentrations of OP50 under vigorous shaking for four to five days, until the majority of the cultured worms were adults. The worm cultures were then chilled on ice, pelleted by centrifugation, washed with M9 (3 g KH_2_PO_4_, 6 g Na_2_HPO_4_, 5 g NaCl, 1 ml 1 M MgSO_4_, H_2_O to 1l). Lysis buffer (10mM tris-HCl, 150 mM NaCl, 5 mM EDTA, 0,5% NP40, 1x PI) and 0.5 mm glass beads (Biospec, USA), were added. Lysis was performed using a FastPrep-24 5G (MPbio, USA) with 5,5 m/s for 120s with 180s break repeated for 3 times at 4°C.

For co-immunoprecipitation, the worm lysates were incubated with GFP-Trap magnetic beads (Chromotek, Germany) in end-to-end rotation at 4 °C for 1 hour. The beads were then separated from the total lysate and washed in lysis buffer and eluted from the beads using SDS sample buffer (Biorad, USA). The eluates were sent for analysis at the Proteomics Core Facility at the Sahlgrenska Academy, University of Gothenburg, Sweden, for liquid chromatography mass spectrometry (LC-MS) analysis.

### Quantification and statistical analysis

All values are shown as mean ± standard error of the mean (SEM).

#### EM image Analysis

Tissue areas were manually traced in IMOD^69^ and total contour areas were used for normalisation. Gold particles were annotated manually in IMOD and assigned to different compartments. Gold particle clusters were identified using the DBSCAN algorithm with a physical distance threshold of 70 nm and a minimum cluster size of three particles. Each cluster was classified according to the dominant tissue type of its constituent particles. Gold particle density and cluster density were calculated as counts per µm² of tissue.

#### Light microscopy

The raw images were processed using ZEN software (Zeiss, Germany) before maximum intensity z-projections were generated in Fiji/ImageJ^70^ for analysis. Semi-automated puncta quantification was carried out using an ImageJ macro^35,36^. For each animal, a single region of interest (ROI) covering the nerve ring and associated cell bodies was selected. The macro quantified the total number of puncta and puncta density (puncta per 10 µm²). To ensure consistency, images were cropped as necessary to match the magnification across the dataset. The data was exported to Prism10 (GraphPad Software, MA, USA) and statistically compared using a paired t-test.

#### Behavioural analysis

Chemotaxis and learning index was calculated as described before^33^. At the end of each assay, worm distributions were counted, and a choice index (CI = (n_test_-n_control_)/n_total_) was calculated. The learning index was defined as the difference between the choice index of untrained and trained animals.

#### Immunoprecipitation analysis

Raw data files contained quantitative metrics for each identified protein, including peptide counts, Peptide Spectrum Matches (PSMs), and unique peptide counts. Gene names were extracted from UniProt’s protein description using pattern matching for the “GN=” identifier. To prevent artificial inflation of fold-change values, duplicate protein entries were identified based on matching gene names and subsequently aggregated. Because the experiment was performed with a single replicate per condition, traditional significance testing was not applied. Instead, protein abundance differences were assessed using fold-change analysis, calculated as log₂FC = log₂((PSM_LGC50Δ363-379}+ 1) / (PSM_LGC50 + 1)), where pseudocounts (+1) were added to avoid division by zero. Proteins were categorised into 11 functional groups based on keyword matching. Keyword searches incorporated both domain-specific and functional terms such as “kinase,” “channel,” “receptor,” “transporter,” “ribosomal,” “cytoskeleton,” and “metabolism.” All visualisations were generated in R using the ggplot2 package. Volcano plots were created with log₂ fold change on the x-axis and total PSM count (log₁₀ scale) on the y-axis, with proteins color-coded by functional category. Protein abundance comparisons between conditions were visualised using log₁₀-transformed PSM counts, and correlation analyses were performed using both Pearson and Spearman methods to assess data consistency. All analyses were conducted in R (version 4.0 or later) using dplyr for data manipulation, ggplot2 and ggrepel for visualisation, and readxl and writexl for Excel file handling.

## Acknowledgements

The authors gratefully acknowledge Andrea Cellini for AlphaFold modelling, Lara Preüssner for help with cloning design and help with generating and maintaining strains and other past and present members of the Morud Lekholm lab for helpful discussions. We would also like to acknowledge Iris Hardege for critical discussions and valuable comments during manuscript writing. We would like to acknowledge the Centre for Cellular Imaging at the University of Gothenburg, Sweden, and the National Microscopy Infrastructure, NMI (NMI), for providing imaging facilities. We would also like to thank the Proteomics Core Facility at the University of Gothenburg, Sweden, for assisting with mass spectrometry analysis. This work was supported by grants from the Swedish Research Council (2022-03951), Swedish Foundation for Strategic Research (FFL21-0166), Tore Nilssons stiftelse, Åke Wibergs stiftelse (M22-0215, M21-0229), Magnus Bergvalls Stiftelse to J.M, the Swedish Research Council (2019-04004) and Cancer Foundation (21 1865 Pj01H) to J.L.H and Lundgrenska stiftelserna to L.C.

## Author contribution

Conceptualisation: J.M., L.C., J.L.H., data curation: L.C., methodology: J.M., J.L.H., L.C., D.P., investigation: L.C., E.A., D.Z., formal analysis: L.C., J.M., visualisation: L.C., J.M., funding acquisition: J.M., J.L.H., project administration: J.M., supervision: J.M., validation: J.M., J.L.H., resources: J.M., J.L.H., writing – original draft: J.M., writing – review and editing: J.M., L.C., D.Z., E.A., J.L.H.

## Data and code availability

All experimental data, including results of imaging, behavioural and biochemical experiments, will be shared by the lead contact upon request. Any additional information required to re-analyse the data reported in this paper is available from the lead contact upon request.

## Supporting Material with Legends

**Table 1.**
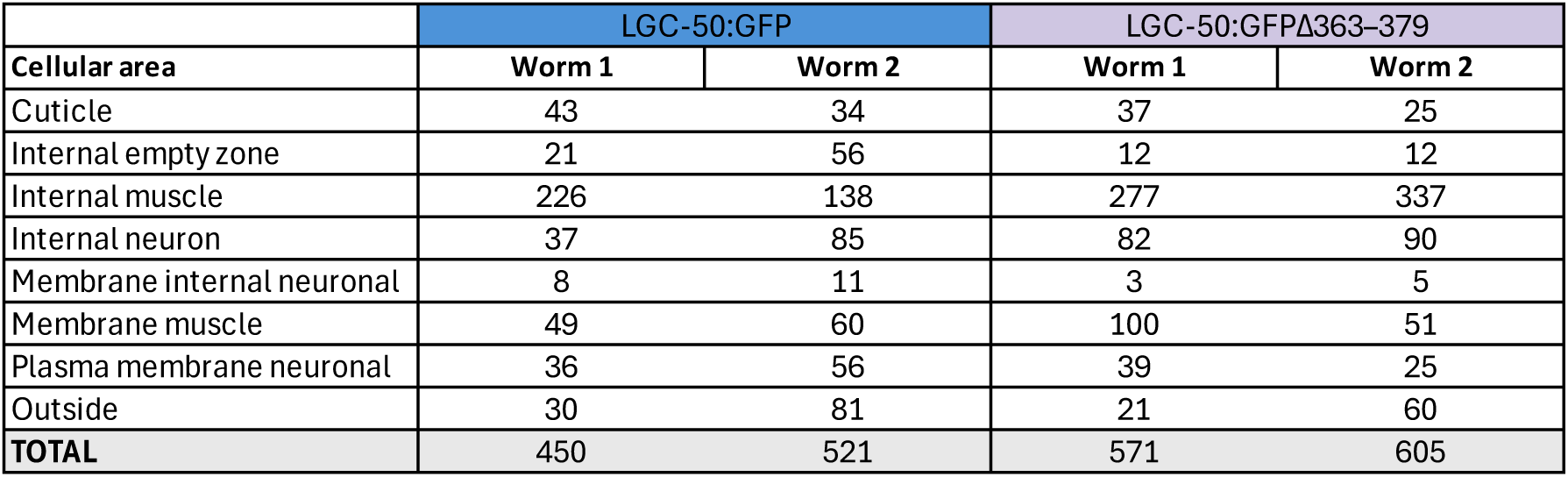
Total number of gold particles detected in different cellular areas of GFP-tagged full-length LGC-50 and 363-379 truncated receptor.

| Cellular area | LGC-50:GFP | | LGC-50:GFPΔ363-379 | |
| --- | --- | --- | --- | --- |
|  | Worm 1 | Worm 2 | Worm 1 | Worm 2 |
| Cuticle | 43 | 34 | 37 | 25 |
| Internal empty zone | 21 | 56 | 12 | 12 |
| Internal muscle | 226 | 138 | 277 | 337 |
| Internal neuron | 37 | 85 | 82 | 90 |
| Membrane internal neuronal | 8 | 11 | 3 | 5 |
| Membrane muscle | 49 | 60 | 100 | 51 |
| Plasma membrane neuronal | 36 | 56 | 39 | 25 |
| Outside | 30 | 81 | 21 | 60 |
| **TOTAL** | 450 | 521 | 571 | 605 |

**Supporting Figure 1.**
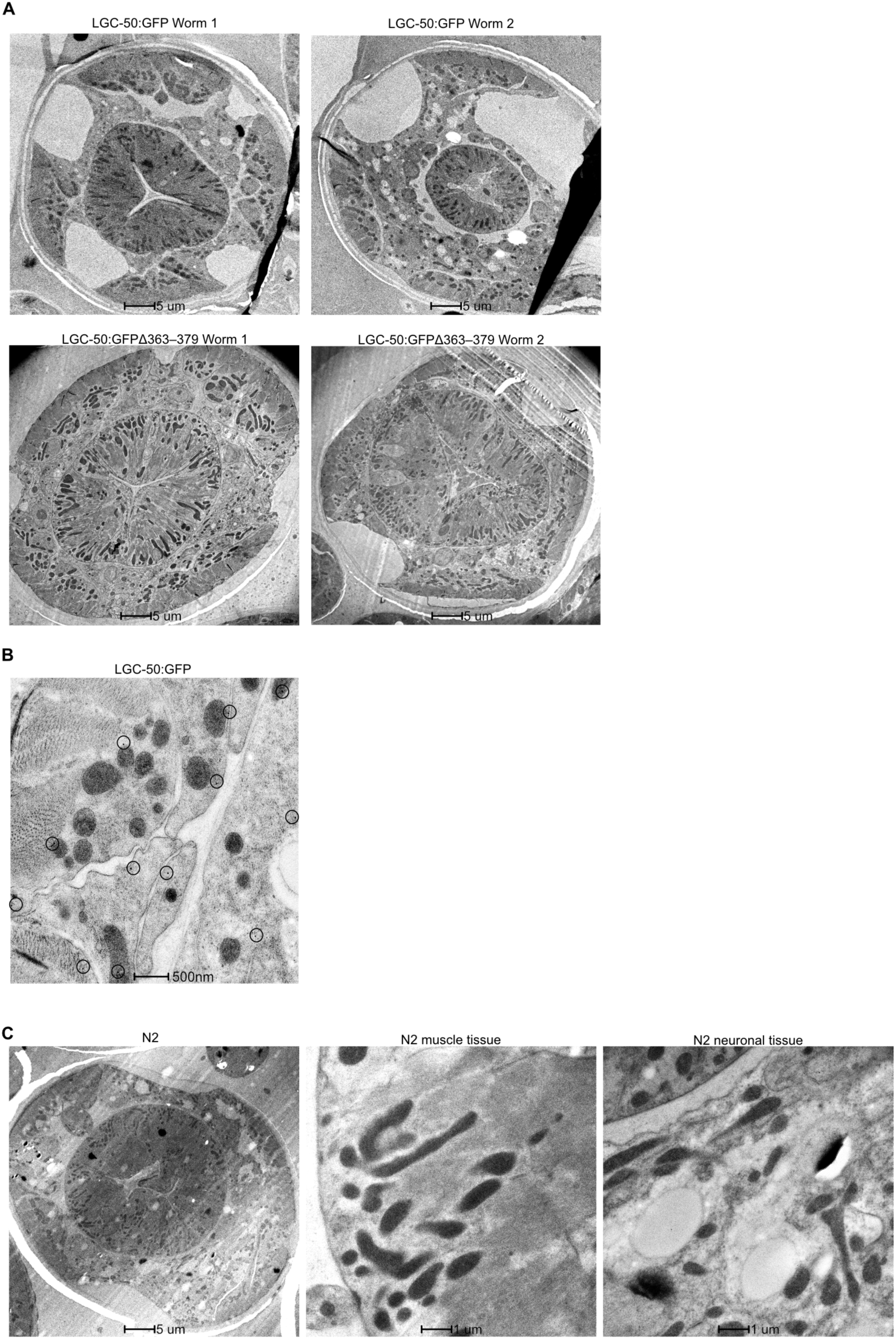
A. Overview of worm sections where high-resolution micrographs were acquired. B. Micrograph example of LGC-50::GFP with immuno-gold labelling. C. EM images from N2 worms after gold labelling, at different magnifications and cellular areas. Very little to no gold particles were found in these sections.

**Supporting Figure 2.**
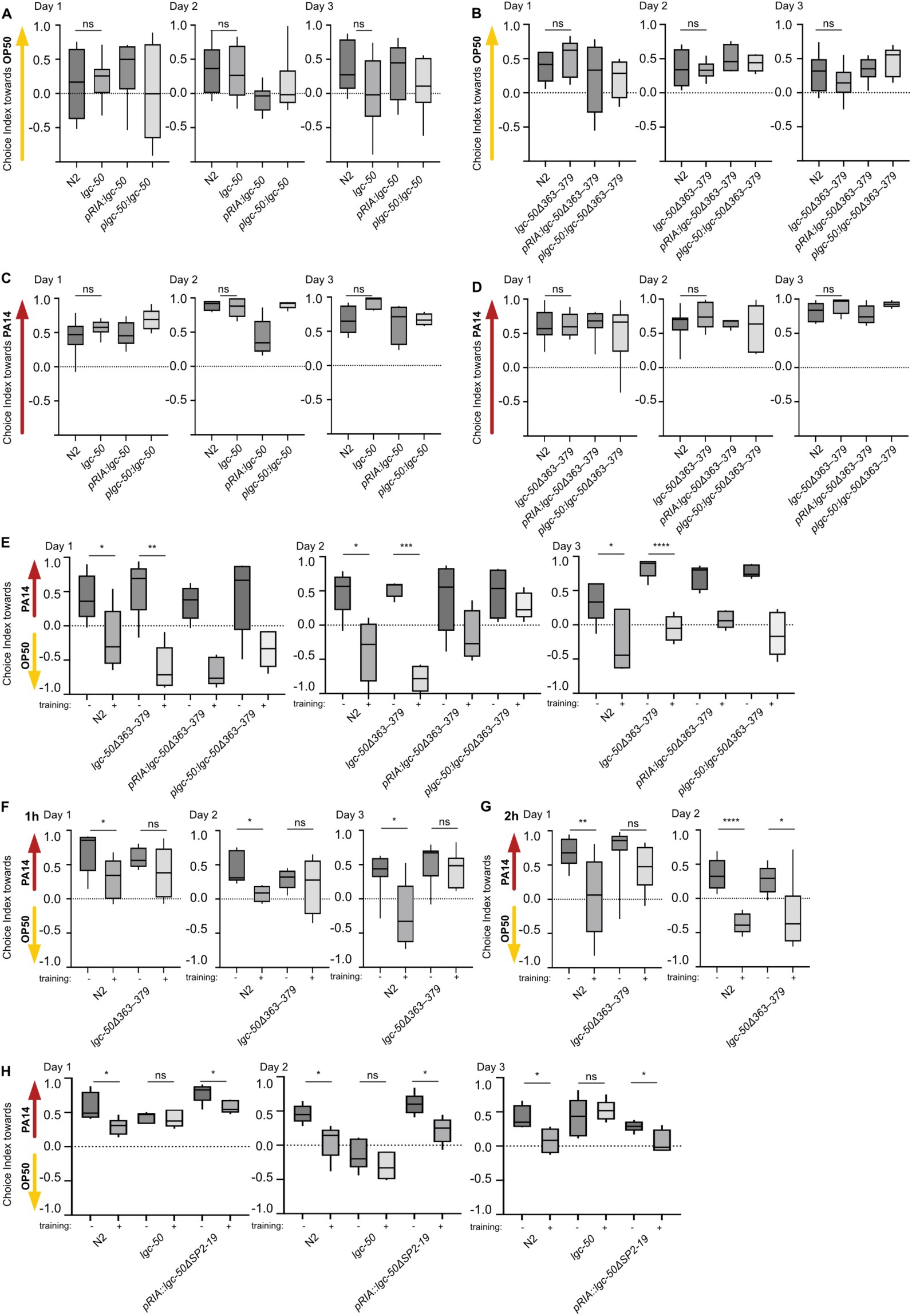
All behavioural tests were done over three independent days to confirm that the day-to-day variability was low and that tests conditions were reproducible. A-B. Choice index calculated for chemotaxis towards OP50 over three different days of experiments. C-D. Choice index calculated for chemotaxis towards PA14 over three different days of experiments. E. Data from three different days for two-choice aversive learning test. F-G. Data from three different days for two-choice aversive learning test with a 1- or 2-hour incubation time after training on PA14. H. Data from three different days for two-choice aversive learning test in signal peptide truncated lgc-50. *P < 0.05. ***P < 0.001, ****P < 0.0001, ns, no significant difference, calculated by Welch’s t-test, data are shown as mean ± SEM.

**Supporting Figure 3.**
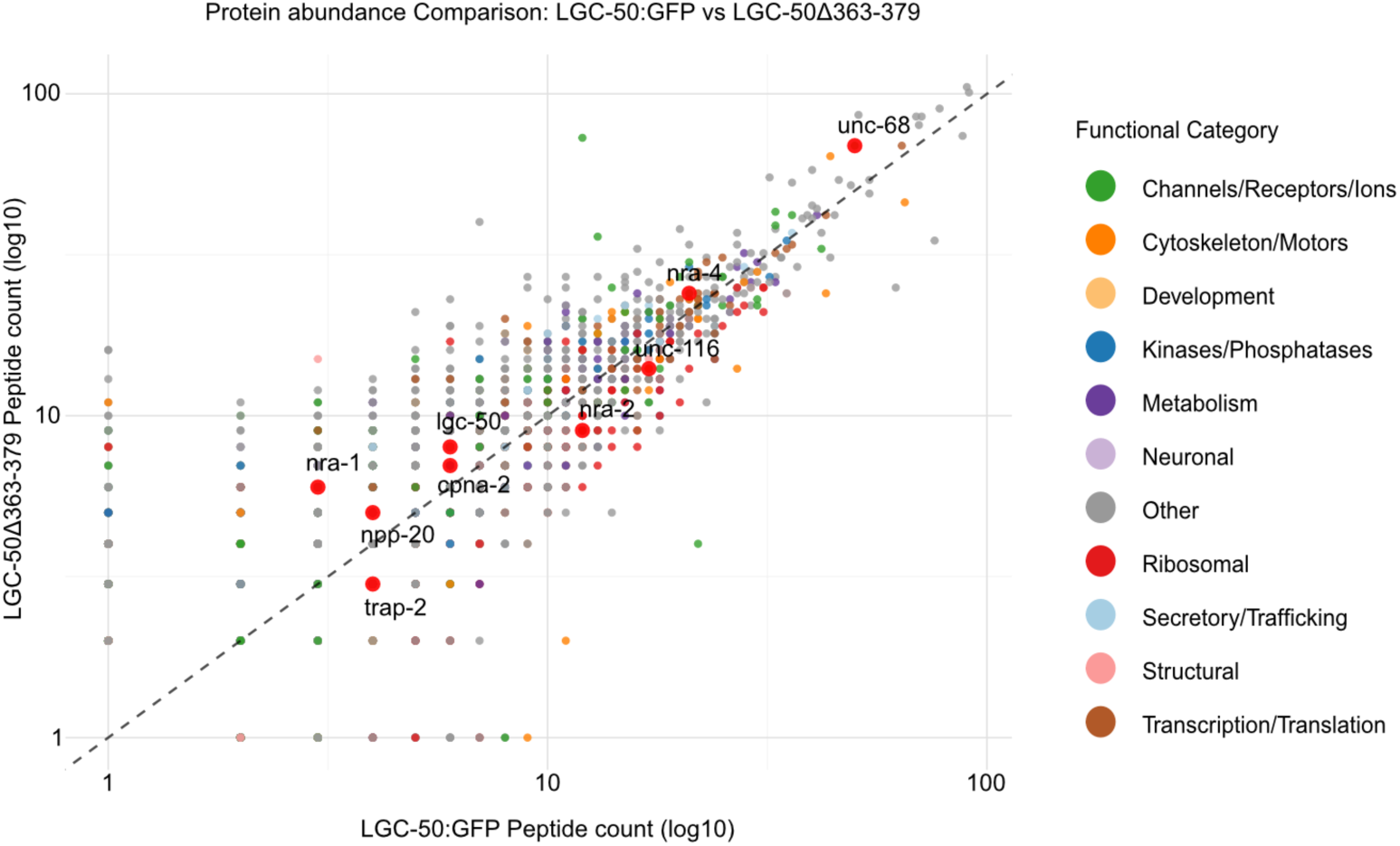
Protein abundance measured by LC-MS after immunoprecipitation of full-length LGC-50::GFP and LGC-50::GFPΔ363-379.

